# LimROTS: A Hybrid Method Integrating Empirical Bayes and Reproducibility-Optimized Statistics for Robust Differential Expression Analysis

**DOI:** 10.1101/2025.02.06.636801

**Authors:** Ali Mostafa Anwar, Akewak Jeba, Leo Lahti, Eleanor Coffey

**Affiliations:** Turku Bioscience Centre, University of Turku and Åbo Akademi University, 20520 Turku, Finland; Department of Computing, University of Turku, Turku, Finland

## Abstract

**Motivation:** Differential expression analysis plays a vital role in omics research enabling precise identification of features that associate with different phenotypes. This process is critical for uncovering biological differences between conditions, such as disease versus healthy states. In proteomics, several statistical methods have been used, ranging from simple t-tests to more advanced methods like limma and ROTS. However, a flexible method for reproducibility-optimized statistics tailored for clinical omics data has been lacking.

**Results:** In this study, we developed LimROTS, a hybrid method integrating the linear model and empirical Bayes method from the limma framework with the Reproducibility-Optimized Statistics from ROTS, to create a novel moderated ranking statistic, for robust and flexible analysis of proteomics data. We validated its performance using twenty-one proteomics gold standard spike-in datasets with different protein mixtures, MS instruments, and techniques for benchmarking. This hybrid approach improves accuracy and reproducibility of complex proteomics data, making LimROTS a powerful tool for high-dimensional omics data analysis.

**Availability and Implementation:** LimROTS has been implemented as an R/Bioconductor package, available at https://bioconductor.org/packages/LimROTS/. Additionally, the code used in this study is available in GitHub repository https://github.com/AliYoussef96/LimROTSmanuscript

**Graphical abstract:** 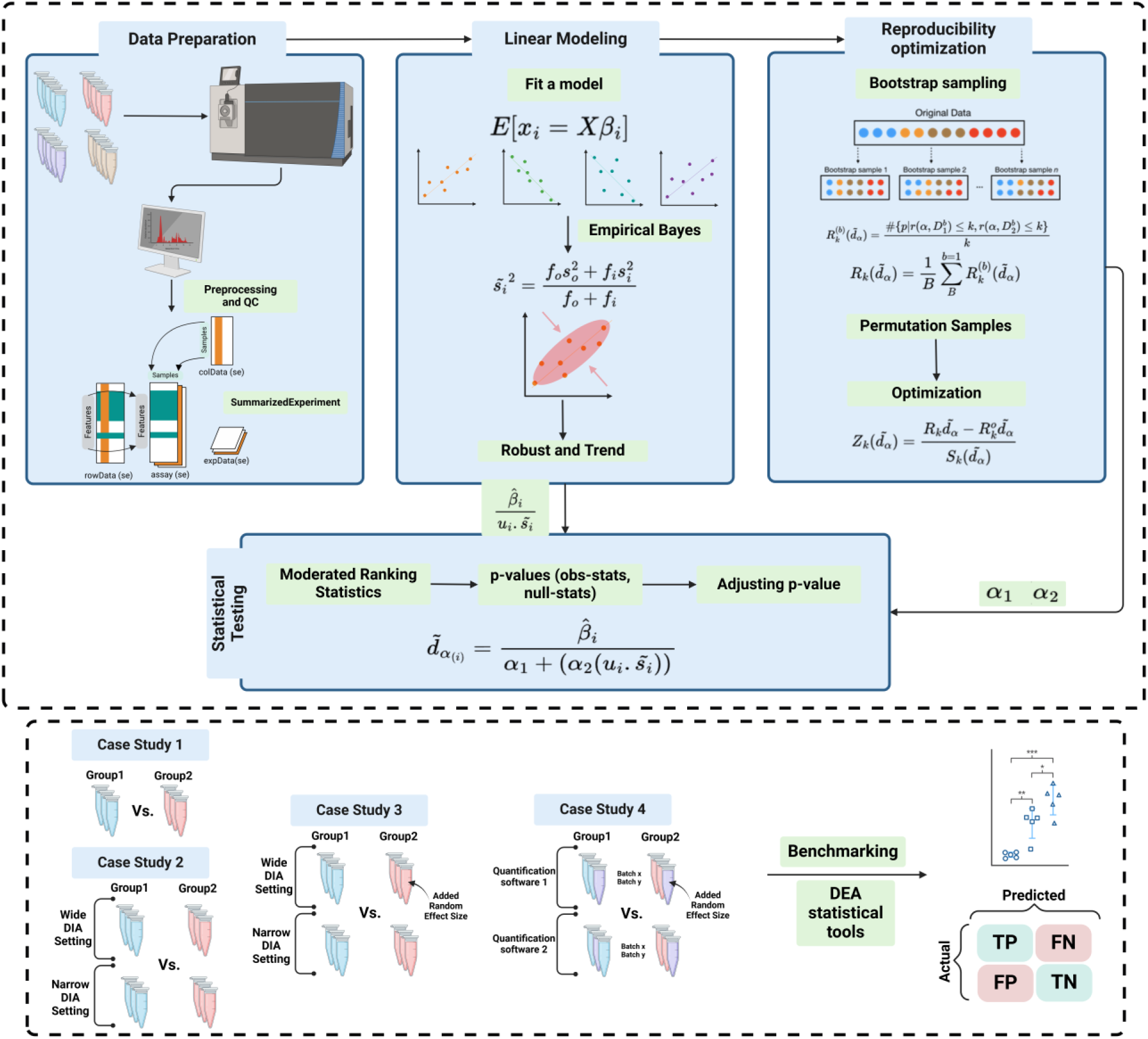

## Introduction

Advances in mass spectrometry technologies and data processing methods have expanded the depth of proteomic detail attainable from biospecimens. These large datasets encode complex biological information that can help to advance precision medicine (Deng et al., 2025; He et al., 2024). The ability to detect phenotype-specific protein changes not only enhances our understanding of underlying biological mechanisms, it also holds promise for the discovery of novel biomarkers for early disease diagnosis, prognosis, and monitoring. Moreover, it aids the identification of potential drug targets for the development of new therapeutic strategies (Meissner et al., 2022; Niu et al., 2022).

One statistical tool that has recently gained attention for its high performance in proteomics data analysis is ROTS (Reproducibility-Optimized Test Statistic). ROTS is a statistical method specifically designed for differential expression analysis (DEA) of microarray RNA data, including applications in proteomics (Dowell et al., 2021; Elo et al., 2009; Peng et al., 2024; Suomi et al., 2017). It identifies differentially expressed proteins (DEPs) by optimizing the test statistic based on reproducibility across bootstrap datasets. Instead of relying on traditional fixed test statistics, ROTS searches for the optimal combination of statistical parameters that maximize the reproducibility of significant results. This data-driven approach improves the detection of true biological signals while controlling for false discoveries (Suomi et al., 2017), making ROTS a robust tool for analyzing complex proteomics datasets.

A highly utilized tool that has been developed for microarray and RNA-Seq data is limma (Ritchie et al., 2015). It applies linear models to the data which help to consider multiple variables and complex experimental designs. limma applies empirical Bayes methods to improve statistical power and control for false discoveries, making it particularly effective for high-dimensional data like proteomics (Peng et al., 2024; van Ooijen et al., 2017). By combining robust statistical techniques with flexible model designs, limma has become a cornerstone tool for identifying DEPs in a variety of biological contexts (van Ooijen et al., 2017).

Although ROTS has shown superior performance in proteomics benchmarking studies (Dowell et al., 2021; Peng et al., 2024), its framework lacks flexibility for complex clinical proteomics datasets, as it cannot account for covariance in the data. For example, the effect of covariates such as age and gender on the data outcome, and technical variations such as batch effects, need to be taken into consideration (Loo et al., 2024; Yan et al., 2024). Omics studies involving DEA have shown that the empirical Bayes method provides a robust approach when implemented within the limma framework (Ritchie et al., 2015). In this study, we developed LimROTS, a hybrid method integrating the linear model and the empirical Bayes method from the limma framework with the Reproducibility-Optimized Statistics from ROTS, by introducing a new moderated ranking statistic, for robust and flexible analysis of proteomics data. To test LimROTS, 21 gold standard spike-in datasets (Peng et al., 2024) were utilized and compared to limma, ROTS, ANOVA, t-test, and SAM. Furthermore, we assessed LimROTS, limma, and ROTS in a real-word experiment using Alzheimer’s disease (AD) samples from the University of Pennsylvania School of Medicine Brain Bank and the Baltimore Coroner’s Office (Johnson et al., 2020).

## Materials and Methods

### Integrating Linear Model and Empirical Bayes with Reproducibility-Optimized Statistics

LimROTS integrates the statistical principles of linear model and empirical Bayes as implemented in limma (Ritchie et al., 2015) and extends that as an optimization problem, solved by the reproducibility optimization statistics (Suomi et al., 2017). Therefore, in principle, LimROTS inherits part of the mathematical rationale from both methods, with a new moderated ranking statistic.

The new moderated ranking statistic 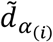 introduced in LimROTS can be represented as:

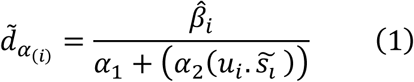

where, 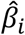 is the effect (coefficient) due to the experiment conditions, *α*_1_ and *α*_2_ are reproducibility optimized parameters, *u*_*i*_ is the un-scaled standard deviation, and 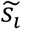 is the posterior residual standard deviation.

To calculate 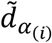, first, a feature-wise linear model is fitted which allow to handle complex experimental design, this can be represented as:

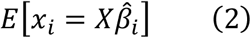

where, *x*_*i*_ is the observed expression value of a feature, *X* is the design matrix of an experiment, and 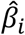 is the effect (coefficient) trying to be estimated by the linear model, in other words; how much expression changes due to the experiment conditions.

Using parametric empirical Bayes technique to borrow information between features in a dynamic way, the *s̃_i_* which is the posterior residual standard deviation is used to adjust the variance to account for uncertainty in the features measurements and combines it with prior information estimated from the data, leading to empirical Bayes variance shrinkage, which can be calculated as;

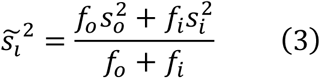

where, *f*_*o*_ is the prior degree of freedom (a global parameter) calculated from the whole dataset, 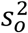 is the prior variance (a global parameter) calculated from the whole dataset, *f*_*i*_ is the residual degree of freedom calculated for each feature (a feature specific parameter), and 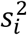 is the observed residual variance for feature *i* (a feature specific parameter).

Equation three allows more reliable estimate of variance for each feature, as well as more statistical confidence in the estimates, because the final variance estimate for each feature (*s̃_i_*) is a balance between two things: the feature specific variance, and the global variance across all the features in the experiment. In other words, each feature specific variance is shrinkage to a common variance estimated from all the features in the dataset.

The optimization parameters *α*_1_and *α*_2_are estimated by maximizing the reproducibility by z-type statistics 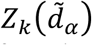. To calculate the 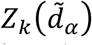, the average reproducibility score 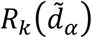 has to be calculated from bootstrap datasets as:

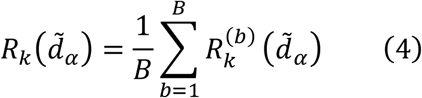

where, for each pair of bootstrap datasets 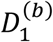 and 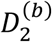 the reproducibility 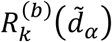 can be computed as:

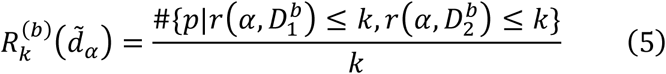

where, {*p*|… } represents the set of features that satisfy the condition of being less than or equal to *k* (*k* equal to cutoff of considering a feature as a top rank feature) in both 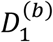 and 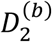, 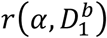 is the rank of feature *p* in the dataset 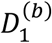 with respect to the statistic 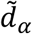 and the same for 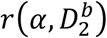, then scaled by the *k*.

Finally, the 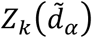 can be estimated by:

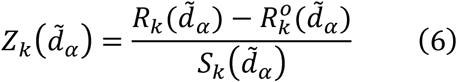

The standard deviation 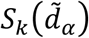 can be computed by square root the variance, calculated as;

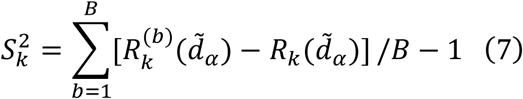

where, *B* is the number of bootstrapping datasets. The 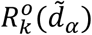 is the null reproducibility estimate of 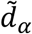 in random (permutated) data. Finally, to avoid the need of pre-specified *k*, LimROTS maximize the 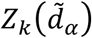 over a lattice of (*α*, *k*)-pairs.

### Datasets used for workflow benchmarking

Table 1 shows twenty-one gold standard spike-in datasets used in this study to test LimROTS performance against other tools. Datasets were adapted from the published benchmark study (Peng et al., 2024). For DIA quantification, Spectronaut (Bruderer et al., 2017) and DIANN (Demichev et al., 2019) were used, with directLFQ (Ammar et al., 2023) as expression matrix, and missing value were imputed by sequential imputation using impSeq function in R (Verboven et al., 2007) or by left-censored imputation using MinDet function (Lazar & Burger, 2022). For DDA, MaxQuant (Prianichnikov et al., 2020) and FragPipe (Kong et al., 2017), with directLFQ as expression matrix, and missing value were imputed by sequential imputation using impSeq and seqKNNimp (Aleš & Marjan, 2023) functions. Sample size ranged from 6 to 35 with conditions from 2 to 9, which means a dataset may have more than one contrast; therefore, all the contrasts for datasets have been considered in the comparison. The quantified features in these datasets extended from 1067 to 11310. In Case Study 1, all the datasets have been analyzed in their original form (after preprocessing), with 66 comparisons in total. To add complexity to the experimental design to mimic the bias that may exist in a real-word biological experiment rather than gold standard spike-in datasets. In Case Study 2, we merged both HEof_w60 and HEof_n600 datasets, as they were identical samples analyzed twice with two different quantification settings (wide and narrow DIA settings), simulating a batch effect. Nonetheless, this is still simpler than an actual batch effect, as these samples from both wide and narrow DIA sittings are the same.

**Table 1:**
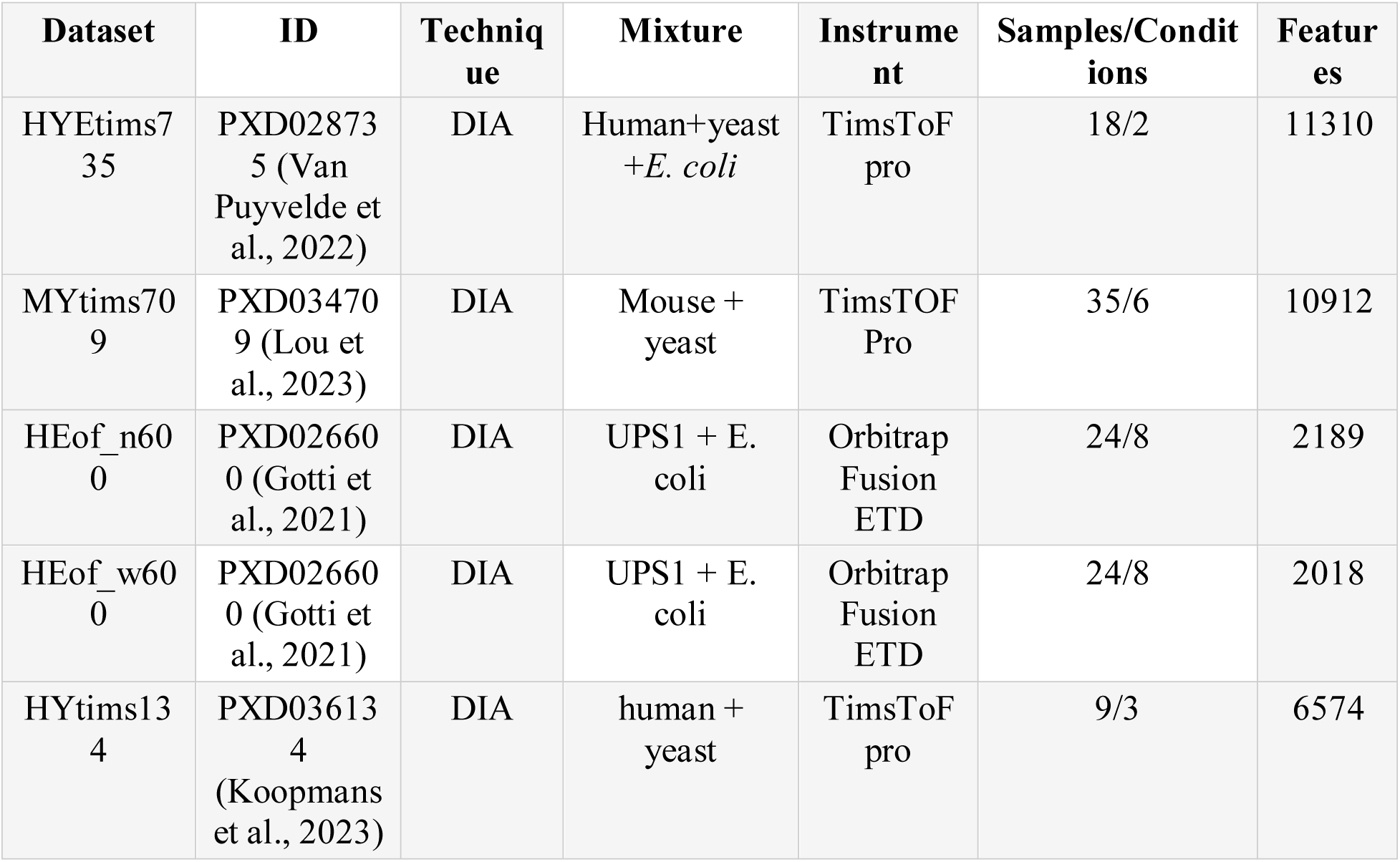
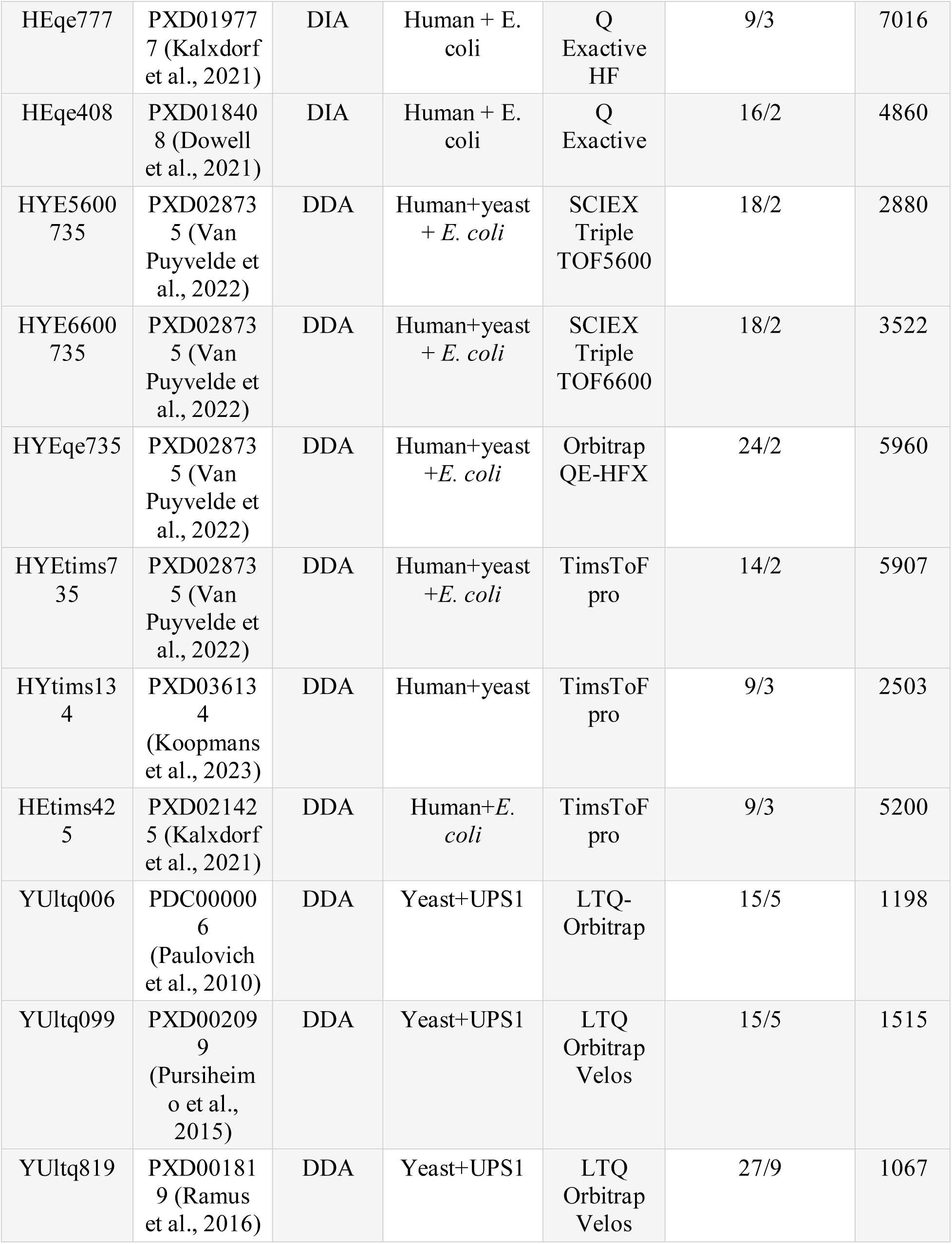
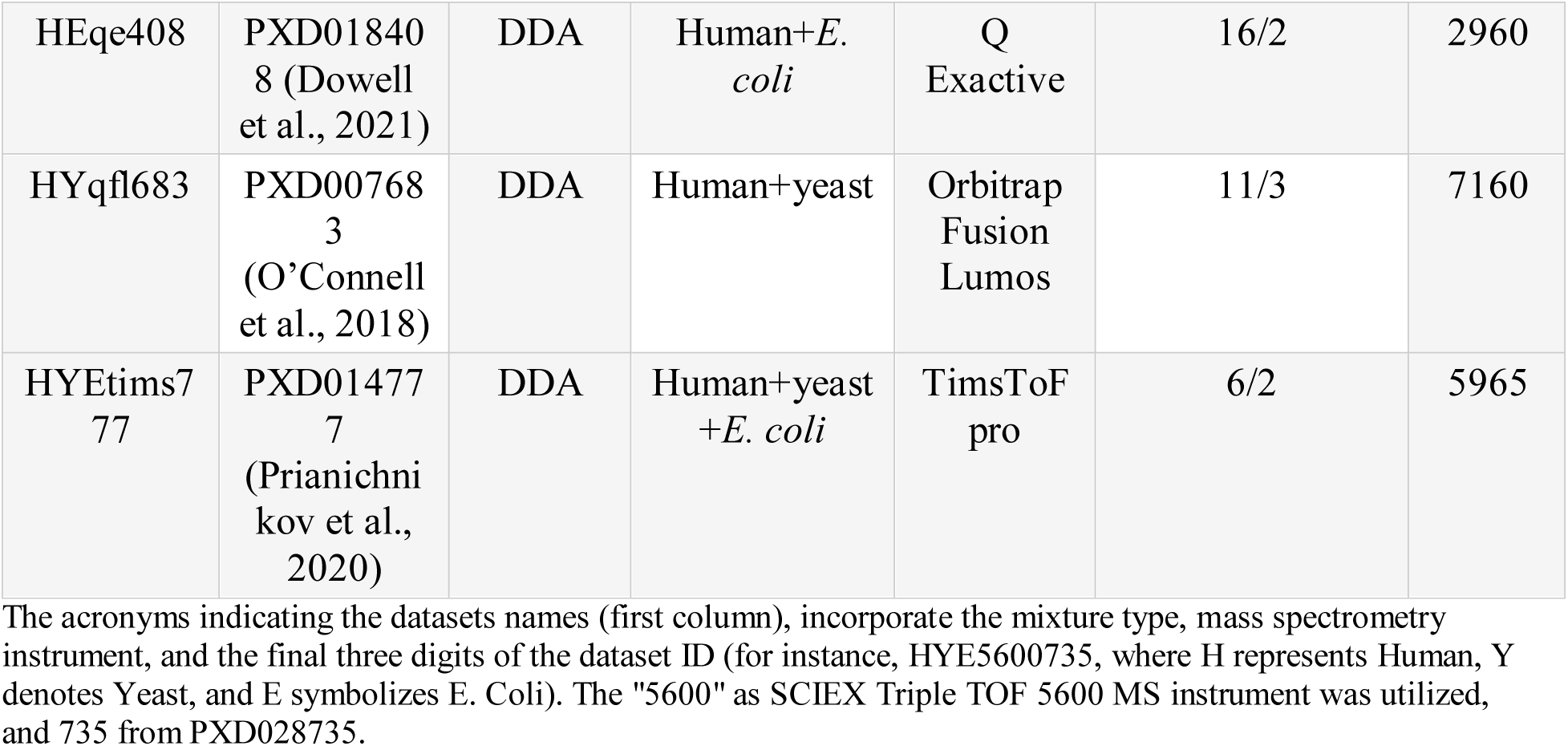
Datasets used for workflow benchmarking.

In Case Study 3, employed the same sample mixture used in Case Study 2. However, in Case Study 3, we added a random effect size ranging from 5 to 20 to 500 random E. coli proteins (which were expected to exhibit no variation between the contrasts) only in the wide DIA samples of one contrast. This will produce wide DIA samples with batch effect, that different from in the narrow DIA samples, introducing bias that may be encountered in actual biological experiments.

In Case Study 4, the dataset was employed from (Gotti et al., 2022). In this instance, we used alternative methodology compared to case study 3. We combined Samples with different UPS1 spiked-in concentration ranging from 0.1 to 50 fmol per microgram of E. coli proteins, to form two groups: one with low UPS1 concentration (from 0.1 fmol to 2.5 fmol) and another with high UPS1 concentration (from 5 fmol to 50 fmol), rather than treating each concentration as a separate group. Subsequently we assigned two toy batches labels to each group of samples with an unbalanced ratio, then we added an effect size of 10, to 100 random E. coli proteins to all samples in one of the toy batches. Furthermore, the same samples in both concentration groups were quantified twice using Spectronaut (Bruderer et al., 2017) and ScaffoldDIA (Searle et al., 2018), both were considered in the same dataset.

UPenn cohorts were performed on a Q-Exactive Plus mass spectrometer essentially as described in (Seyfried et al., 2017) Then, using the MaxQuant protein groups results, we set the razor unique peptides number to be two or more, proteins groups showed more than 50% of missing value were removed, impSeq function was used for imputing the rest of the missing values, and ComBat from SVA package (Leek et al., 2024) was used to correct for the batch effect. Finally, we performed DEA between the AD and the Control group, using LimROTS, limma and ROTS. The parameters and imputation methods used in preprocessing all the datasets in this study, were selected based on recommendations and results from (Peng et al., 2024).

### Significant cutoffs and Performance evaluation metrics

Throughout this study, significant proteins were identified with both 0.01 and/or 0.05 adjusted p-values, additionally, since in all the dataset contrasts was compared in a way that the high concentration samples were used as a target group, therefore, only proteins showed upregulation in term of fold change were considered as true positive (otherwise considered as false positive). Five evaluation metrics were used in this study to evaluates the methods performance. Normalized Matthews correlation coefficient (nMCC), harmonic mean of precision and recall (F1 score), geometric mean of sensitivity and specificity (G-mean), balanced accuracy, and the partial area under the ROC (pAUC) with two FDR cutoffs 0.01 and 0.05. The median of each matrix is utilized to summarize the performance across the datasets in each case study, for each benchmarked method.

## Results

### Case study 1: Evaluating LimROTS performance in gold standard spike-in datasets

In Case study 1, all possible contrasts have been analyzed for each dataset, with an average of 2 contrasts per dataset, corresponding to approximately 66 comparisons overall in DIA and 71 in DDA. When using DIANN as quantification software, with a significance threshold of 5% FDR, LimROTS and ROTS outperformed other methods in terms of nMCC (median values: 0.88 and 0.87, respectively), F1 score (median values: 0.79 and 0.75, respectively), and pAUC 5% (median values: 0.94 and 0.92, respectively) (Figure 1A). However, LimROTS consistently displayed slightly higher values than ROTS across all these evaluation metrics. A similar trend was observed when Spectronaut was used as the quantification software. The results for all evaluation metrics across datasets and tools used in this study are available in Supplementary File 1, Table S1 (A–D). Furthermore, with the DDA dataset using MaxQuant as the quantification software, LimROTS and ROTS remained as the top-performing methods in terms of F1 score and nMCC (Supplementary File 1, Table S1 G–H). Nevertheless, when FragPipe was used as the quantification software, LimROTS and ROTS continued to be the top-performing methods overall; however, performance across all tools was notably lower in terms of F1 score and nMCC (Supplementary File 1, Table S1 E–F).

**Figure 1:**
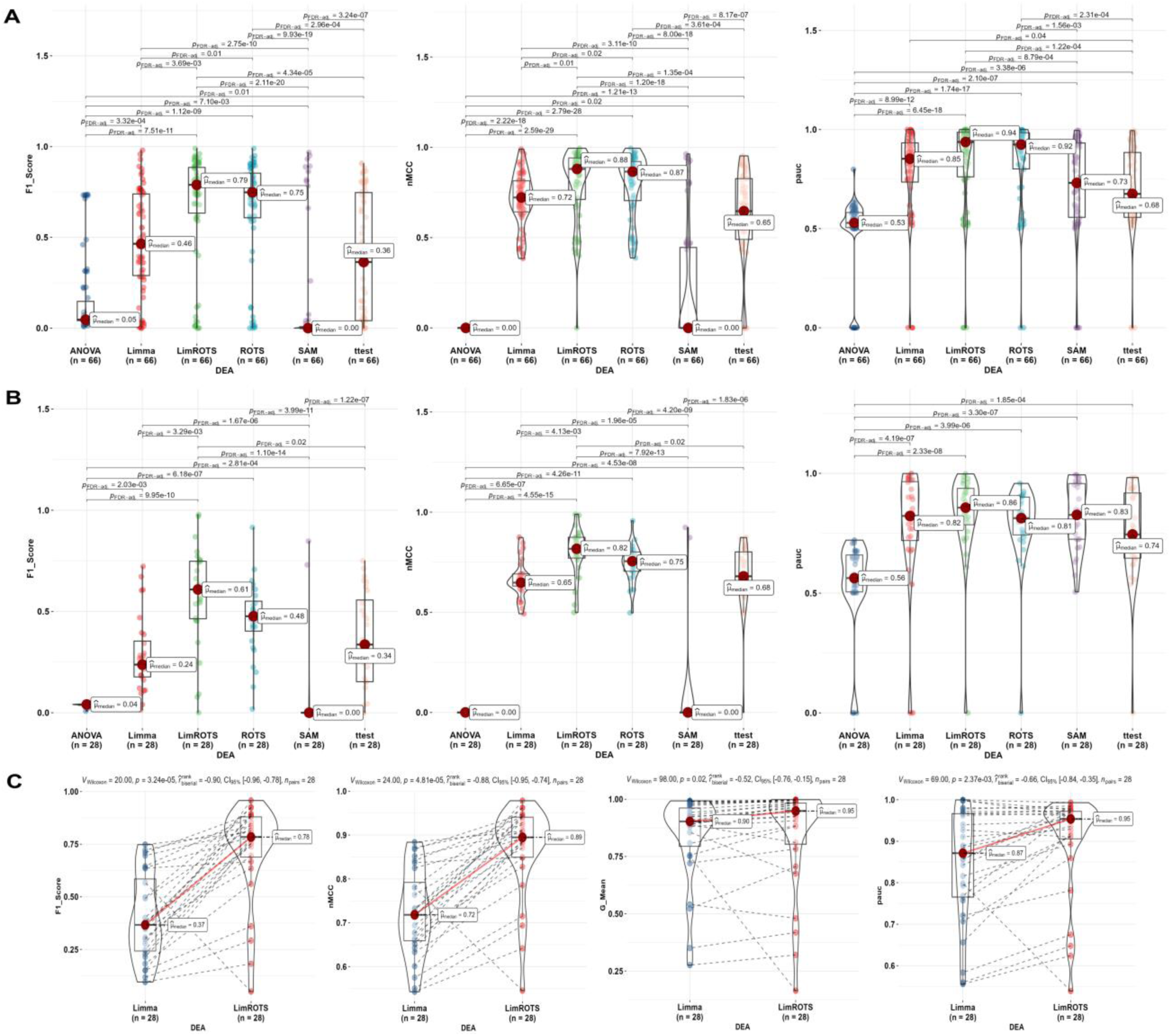
Benchmarking Performance Metrics; F1 score, nMCC, and pAUC5%. A: The benchmarking performances results (F1 score, nMCC, and pAUC5%) boxplots (with violin plot when it possible) for each method from case study 1 (Quantified by DIANN software), with adjusted p-values for the methods showing statistically significant differences. B: The benchmarking performances results (F1 score, nMCC, and pAUC5%) boxplots (with violin plot when it possible) for each method from case study 2 (Quantified by DIANN software), with adjusted p-values for the methods showing statistically significant differences. C: Presents the results of the benchmarking performances (F1 score, nMCC, G-mean and pAUC5%) boxplots (with violin plot when it possible) between LimROTS and limma, from case study 3 (Quantified by Spectronaut software), with adjusted p-values for the methods showing statistically significant differences.

### Case Study 2: Dataset with two different DIA settings simulating a batch effect

The dataset created for this case study is more complex than the previous one, as it focuses on the HEof dataset (PXD026600, (Gotti et al., 2021)), combining both wide DIA and narrow DIA settings using the same samples. Using DIANN as the quantification software, LimROTS showed a notably higher performance in F1 score (median 0.61, with ROTS second at 0.48), nMCC (median 0.82, with ROTS second at 0.75), and pAUC 5% (median 0.86, with ROTS second at 0.81) (Figure 1B). A similar trend was observed when Spectronaut was used as the quantitation software. The results for all evaluation metrics across all datasets and tools used in this case study can be found in Supplementary File 1, Table S2 A–D.

### Case Studies 3 and 4: Evaluating LimROTS and limma in more complex semi-synthetic datasets

In these case studies, we excluded SAM, ROTS, and the t-test, as these methods are not specifically designed to account for covariates in an experiment, which is crucial for obtaining accurate results, as demonstrated in Case Study 2. Additionally, ANOVA was excluded this time due to its overall poor performance. (comprehensive results for all tools are provided in Supplementary File 1, Table S3 A–D.) Therefore, LimROTS was compared against limma in this analysis. In case 3, Using Spectronaut as the quantification software, LimROTS demonstrated significantly higher performance across all evaluation metrics compared to limma. Specifically, LimROTS achieved values of 0.78 (F1 score), 0.89 (nMCC), 0.95 (pAUC 5%), 0.95 (G-mean), while limma achieved values of 0.37, 0.72, 0.87, and 0.9, respectively, as shown in Figure 1C. Similarly, using DIANN as the quantification software, LimROTS outperformed limma in nMCC (0.81 compared to 0.65) and F1 score (0.57 compared to 0.25) (Supplementary File 1, Table S3 A–D).

In Case 4, as shown in Figure 2A, LimROTS notably outperformed limma across all evaluated metrics. Specifically, LimROTS and limma achieved the following scores: F1 score (0.78 compared to 0.15), nMCC (0.90 compared to 0.61), pAUC 5% (0.96 compared to 0.90), G-mean (0.99 compared to 0.81), and balanced accuracy (0.99 compared to 0.83), respectively.

**Figure 2:**
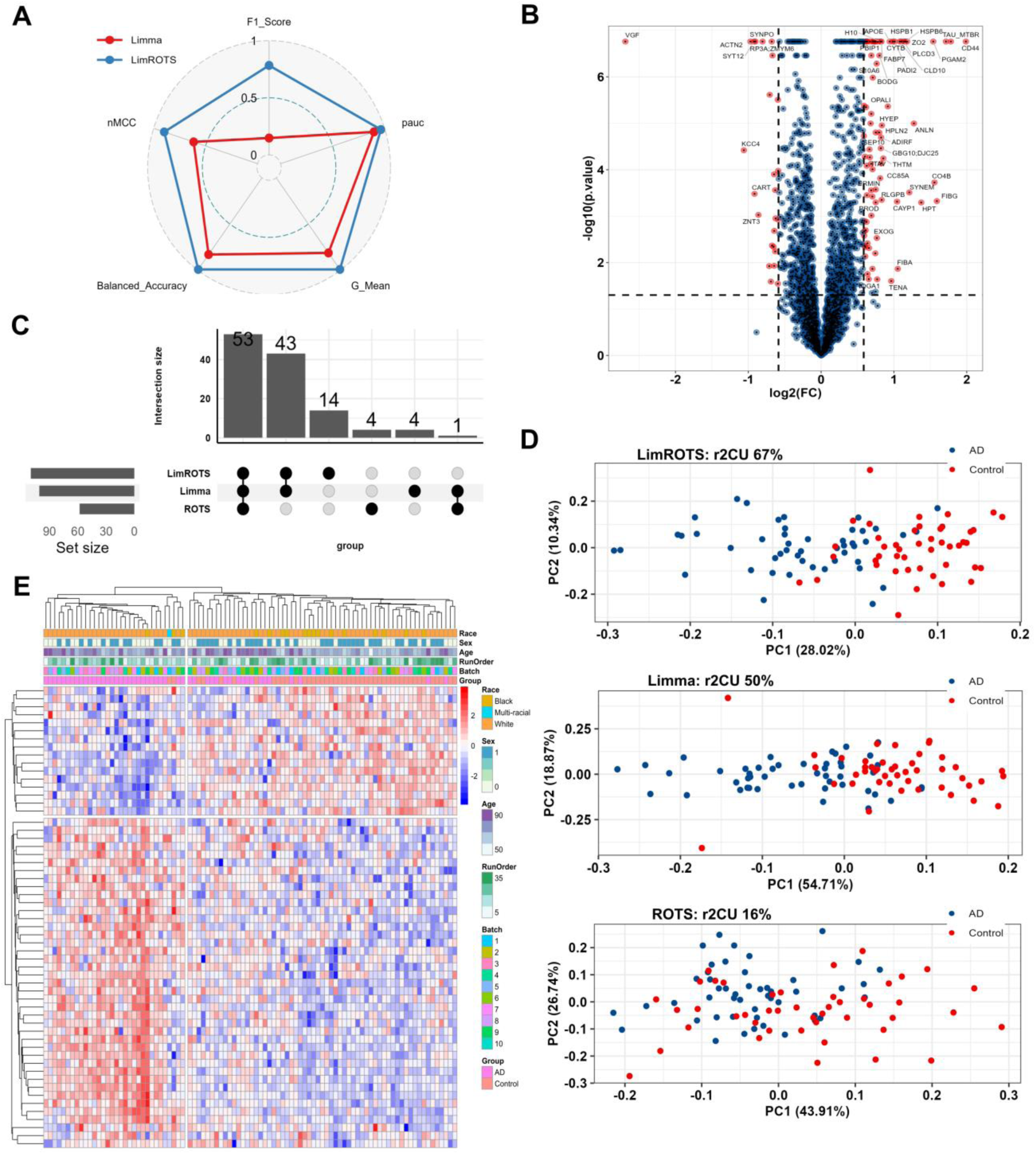
Benchmarking Performance for Case study 4 and UPenn cohort results. A: A spider plot illustrates the performance differences between LimROTS (blue) and limma (red) in case study 4, using five metrics: F1 score, nMCC, pAUC5%, G-mean, and balanced accuracy. B: The volcano plot shows the DEPs (p-value < 0.05 and FC >= log2(1.5)) represented by red dots from LimROTS, annotated with gene symbols. C: UpSet displays the overlap DEPs between the three methods (LimROTS, ROTS, and limma), defined by the cutoffs, (FDR < 0.05 and FC >= log2(1.5)). D: PCA plots for non-overlapped DEPs identified using each method, from LimROTS, limma, and ROTS. Red and blue dots represent the Control and AD groups respectively. Above each PCA, the Cragg and Uhler’s pseudo r-squared (r2CU) from fitting the PC1 with GLM, is displayed. E: Heatmap with a hierarchical clustering to the samples and LimROTS DEPs. Samples were annotated with Race, Sex, Age, Ms Run Order, and Digestion batches.

### UPenn Case study: Performance of DEA tools using real-world clinical proteomics data

In order to evaluate LimROTS performance with real-world clinical data, we used the UPenn cohort. This cohort contains post-mortem brain samples from a range of neurodegenerative diseases. We used only the Alzheimer’s disease (AD) cases compared to healthy controls. DEPs were selected with a cutoff of 5% FDR and log2 1.5-fold change (supplementary file 2, Tables S1-3 and Figure 2B). The UpSet plot (Figure 2C) presents the number of DEPs that overlap and the number that are unique, comparing LimROTS, limma and ROTS. To estimate to what extent the unique proteins from each test represented biological differences in AD Vs Control, a PCA was conducted (Figure 2D). We then fitted a generalized linear model (GLM) to PC1 with diagnostic (binary) status as a response variable (AD Vs. Control), then Cragg and Uhler’s pseudo r-squared was computed (r2CU). Additionally, we calculated the receiver operating characteristic (ROC) curve for the prediction’s probability of each model. LimROTS achieved the highest r2CU (67%) and 0.93 area under the ROC (AUC), followed by limma with r2CU equal to 50% and 0.87 AUC, lastly ROTS achieved only 16% and AUC equal to 0.7 (Figure S1). Next, the DEPs from LimROTS (Figure 2B) were used for enrichment analysis using several databases; DisGeNET, Jensen DISEASES, KEGG, and Human Phenotype Ontology, using Enrichr server (Xie et al., 2021). With significant cutoff < 0.05 p-value (Figure S2). Moreover, a heat-map with a hierarchical clustering, annotating Race, Sex, Age, Ms Run Order, and Digestion batches was drawn (Figure 2E).

## Discussion

In Case study 1, the findings suggest that LimROTS and ROTS offer superior overall performance compared to alternative methods (e.g., limma, t-test, ANOVA, and SAM) when DIANN or Spectronaut is used as the quantification software under a significance threshold of 5% FDR. The high nMCC values, which reflect the overall classification performance, indicate that both LimROTS and ROTS accurately detect most true positive proteins while effectively distinguishing true negatives. The F1 score, a metric that balances precision and recall, was also higher for LimROTS and ROTS, further supporting their superior capability in detecting significant proteins while minimizing false positives and false negatives. Additionally, both methods showed strong performance in pAUC 5%, highlighting their ability to rank the most significant proteins in the top 5%. Furthermore, for DDA-based analysis, particularly when using MaxQuant for quantification, LimROTS and ROTS continued to outperform other methods. These results establish LimROTS as promising candidates for DEA in proteomics DDA and DIA workflows.

The findings from Case Study 2, where the experimental design complexity increased by combining the same set of samples analyzed using both narrow and wide DIA settings, indicate that LimROTS showed significantly higher overall performance compared to all other tools. The highest F1 score, 0.61, was achieved by LimROTS, indicating its superior ability to identify true positives while minimizing both false positives and false negatives. In contrast, the second-highest score of 0.48 was recorded by ROTS, which further suggests that LimROTS is more effective, even in the presence of covariates. Furthermore, LimROTS achieved the highest scores for both nMCC and pAUC 5%, indicating better discriminative power and stronger overall performance in distinguishing between true differential and non-differential proteins. These results can be attributed to the additional advantages LimROTS offers over ROTS, as it flexibly integrates both narrow and wide DIA settings into the linear model as covariates, along with its advantage over limma by optimizing the rank statistics of the proteins.

Comparing both LimROTS and limma in case studies 3 and 4, demonstrate that LimROTS significantly outperforms limma (and all other tools, as shown in the supplementary file 1) in balancing precision and recall when using either Spectronaut or DIANN. LimROTS effectively identifies true positives while minimizing false positives and false negatives, as indicated by its high F1 score. Additionally, the G-mean, which represents the balance between sensitivity (true positive rate) and specificity (true negative rate), was much higher for LimROTS compared to limma. Furthermore, the higher pAUC 5% score of LimROTS highlights its superior ability to rank the top 5% of the most DEPs with greater confidence in its top-ranking results. Moreover, the nMCC score of LimROTS exceeded the performance of limma. Overall, LimROTS consistently outperformed limma when using either Spectronaut or DIANN, which indicates that LimROTS is a more reliable and accurate tool for proteomics data. Case Study 4 particularly features a more complex experimental design than the other case studies, requiring a flexible tool to effectively model covariates, even in unbalanced settings. The results from this semi-synthetic dataset show that LimROTS performs better at accurately identifying DEPs while maintaining a strong balance between sensitivity and specificity.

Finally, by using UPenn cohort, unique proteins that only identified as DEPs by LimROTS (14 proteins) demonstrated the greatest r2CU of 67% and with AUC-ROC 0.93, signifying that the DEPs are attributable to the biological disparities between the studied AD and control groups. In this analysis ROTS showed less significant proteins compared to LimROTS and limma (Figure 2C), which affected its PCA and r2CU dramatically. This can be due to the batch correction method (Combat) used to pre-process the dataset before applying ROTS test, which is essential for the analysis. However, this batch correction step could inadvertently eliminate biological signals (Hui et al., 2024; Phua et al., 2022; Wang & Cao, 2023). In contrast, LimROTS and limma have the advantage of incorporating the batch information as a fixed variable in the model, accounting for it during the DEA. Furthermore, the heatmap illustrated that, the DEPs from LimROTS are not due to technical bias due to digestion batch or MS run batch, for example. Rather, the samples clustered according to diagnostic groups (AD and Control) and not according to technical or unwanted biological variables such as age or sex.

In the UPenn cohort, fourteen new DEPs were identified using LimROTS that were not significant in the original analysis (Johnson et al., 2020). These proteins were regulator complex protein 5 (LTOR5), Th1 cell surface antigen (E9PIM6), chimerin-1 and -2 (CHN1/CHN2), cystine and glycine rich protein 1 (CSRP1), CD9 molecule (A6NN14), adenylate kinase 2 (F8W1A4), nuclear mitotic apparatus protein (NUMA1), metalloprotease 1 (PITRM1), long chain fatty acid coA ligase (ACSBG1), PDZ and LIM domain 5 (PDLIM5), C1orf198, epimerase family protein (SDR39U1) and Arf related protein 2 (ARL15) (supplementary file 2, Table S4). Among the upregulated proteins is chimerin-1, a GTPase-activating protein for p21-Rac. It was previously shown to be regulated at the transcriptome level in late-onset AD (Arzouni et al., 2020). Chimerin was shown to regulate microglial migration to Ab deposits in AD (Chen et al., 2022). PDLIM5, which was also elevated in AD samples, plays a role in neuronal development, synaptic assembly, and dendritic spine formation, and is identical to AD7c-NTP, an AD-related protein, and has been connected with psychiatric disorders and polygenic risk scores for AD (Herrick et al., 2010; Miao et al., 2020). Ragulator complex protein 5 (LAMTOR5) was also shown to be upregulated when analysed with LimROTS. This protein activates mTORC1 which inhibits autophagy and may contribute to protein aggregation in AD (Ma et al., 2021). Proteins downregulated in AD included PITRM1. Loss-of-function in this protein correlates with mitochondrial dysfunction and neurodegeneration (Brunetti et al., 2021).These findings support the earlier associations between these proteins and AD pathogenesis. They were found from this dataset only when LimROTS was utilized.

### Limitations and future perspective

While bootstrapping and the permutation techniques utilized in LimROTs are statistically robust, these techniques are computationally more expensive. We attempted to resolve this issue by incorporating parallel processing. This significantly reduced computational time, but at the cost of increased Random-access memory (RAM) utilization. In future versions, we intend to enhance the workflow to speed up the analysis while minimizing the requirement for substantial computational resources. Initially, we will convert the optimization step into a C++ code and integrate it into R using Rcpp package, this could be expected to increase the speed of this step dramatically. Additionally, the bootstrapping and permutation steps could also be parallelized for simultaneous execution, however, prior to this, we should optimize the code for enhanced speed independent of the parallelization. Furthermore, incorporating additional functionality for data preprocessing and visualization.

### Conclusion

LimROTS consistently outperformed other DEA statistical methods, as limma, ROTS, and SAM, in diverse experimental contexts. It exhibited exceptional performance across essential evaluation measures, irrespective of the quantification approach employed. LimROTS demonstrated efficacy in managing intricate experimental designs, optimizing recall and precision, and addressing biases and imbalanced batches more effectively than alternative techniques. Moreover, it yielded biologically significant outcomes in clinical proteomics data, hence enhancing its credibility. In this study we focused on proteomics data; however, LimROTS may have wider relevance in other omics domains, including transcriptomics and metabolomics, pending additional validation. Ultimately, LimROTS proved to be a more precise, dependable, and adaptable method for DEA in high-dimensional omics research.

## Data availability

The 21 Datasets used in case study 1,2, and 3 are available under the OpDEA resource at http://www.ai4pro.tech:3838 adapted from (Peng et al., 2024), the ProteomeXchange repository IDs and the original study for each dataset are available in table 1. The dataset used in case study 4 are available at ProteomeXchange repository with ID number: PXD026600 (Gotti et al., 2022).

## Code availability

LimROTS has been implemented as an R/Bioconductor package, available at https://bioconductor.org/packages/LimROTS/. Additionally, the code used in this study is available in the GitHub repository: https://github.com/AliYoussef96/LimROTSmanuscript. The package supports the R/Bioconductor SummarizedExperiment data structure (Huber et al., 2015), which enhances interoperability with the vast array of other Bioconductor methods.

## Author contributions statement

AMA designed the algorithm, implemented the software (with input from LL and AJ), and conducted data analysis (with input from LL and EC). AMA wrote the manuscript with input from AJ, LL and EC. All authors read, edited and approved the manuscript. No competing interest is declared.

## Acknowledgments

This project has received funding from, the European Union’s Horizon 2020 research and innovation programme under grant agreement No 952914 / FindingPheno (to LL). The Michael J. Fox Foundation grants MJFF-021587 and MJFF-023714 to EC and the ImmuDocs doctoral program award to AMA.

---

**Supplementary file 1.**

The supplementary file includes tables that display the performance metrics assessed across all case studies for every method. Table S1: Findings from Case Study 1, Table S2: Findings from Case Study 2, Table S3: Findings from Case Study 3, and Table S4: Findings from Case Study 4.

**Supplementary file 2.**

The supplementary file has three tables: Table S1 presents the DEA statistical results from LimROTS (p-value, q-value, and log2 fold change), Table S2 offers equivalent data from ROTS, and Table S3 includes results from limma. Table S4, contains the 14 unique LimROTS DEPs.

