## Supplementary figures and images for "LimROTS: A Hybrid Method Integrating Empirical Bayes and Reproducibility-Optimized Statistics for Robust Differential Expression Analysis"

### Figure S1

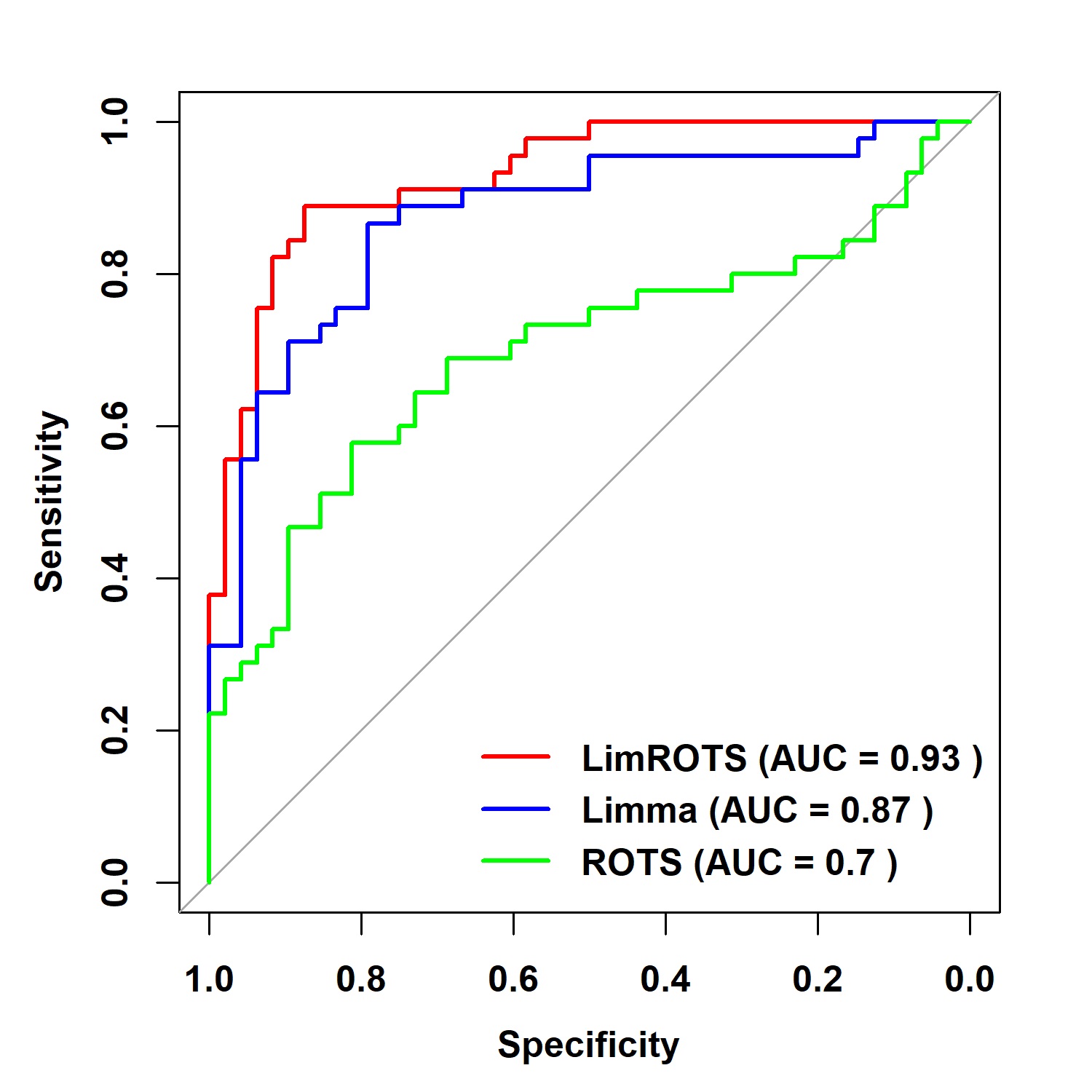

### Figure S2

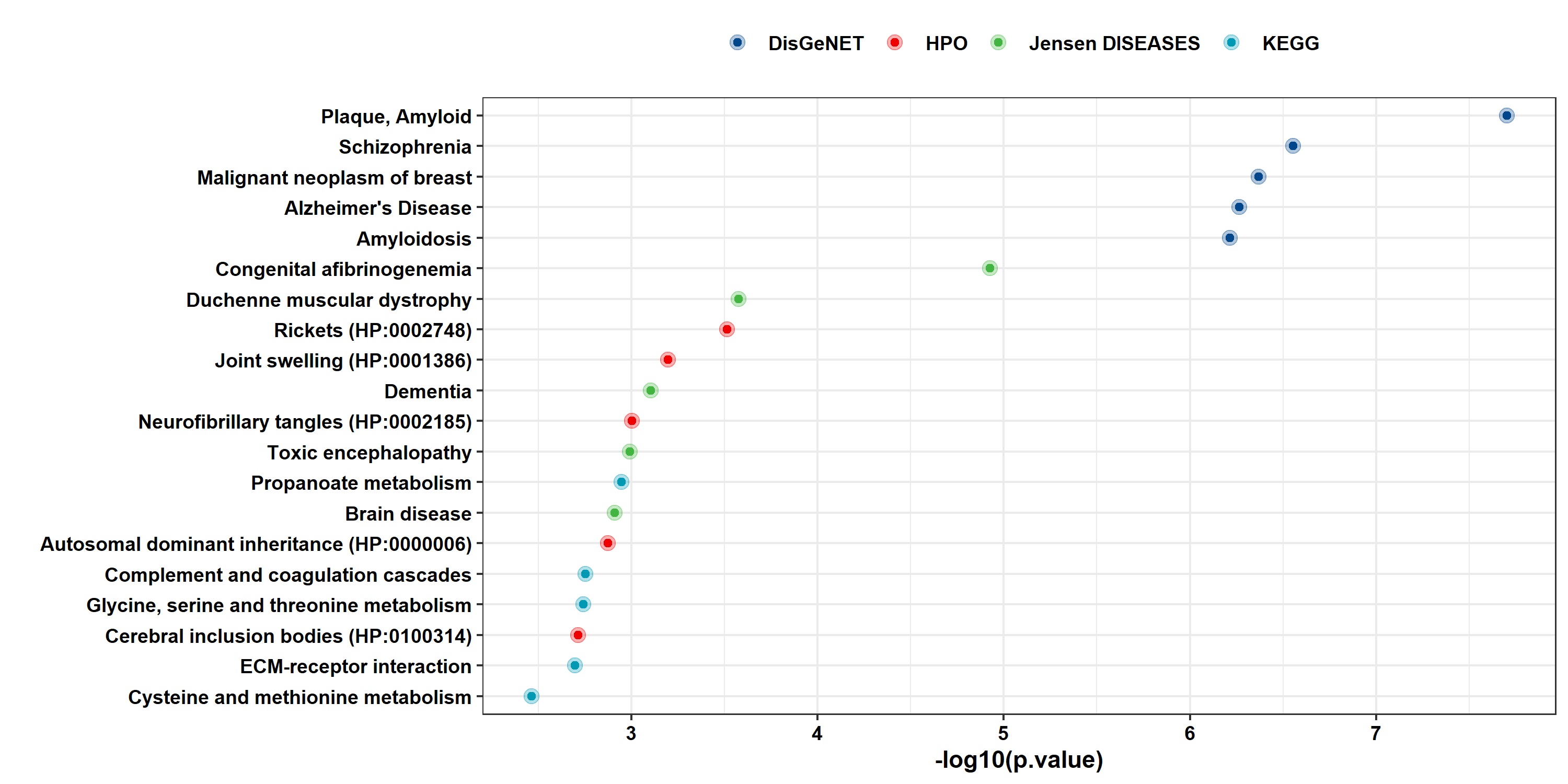
